# Convergent and Distinct Transcriptional Signatures in Suicide, Bipolar Disorder, and Major Depressive Disorder

**DOI:** 10.64898/2026.10.14.759202

**Authors:** Ruchika Mishra, Alexandra M. Wamalwa, Giulia Guasoni, Maura B. Dupont

## Abstract

Globally, a person dies by suicide every 40 seconds, underscoring a major global public health concern. In the United States, suicide is the second leading cause of death among individuals aged 10-34 years. Suicide is a complex, multifactorial phenomenon influenced by genetic and epigenetic factors and is strongly associated with mood disorders, particularly major depressive disorder (MDD) and bipolar disorder (BD). However, the extent to which their molecular signatures converge or diverge across disorders and brain regions remains incompletely understood. To address this gap, we conducted a systematic literature review of suicide, MDD, and BD transcriptomic studies in accordance with PRISMA 2020 guidelines, searching PubMed, Embase, and Web of Science. Eligible studies used human postmortem brain tissue, reported a complete genome-wide transcriptomic list of differentially expressed genes (DEG), and compared suicide decedents with psychiatric non-suicide individuals diagnosed with the same underlying disorder. Regarding BD, studies comparing affected individuals to unaffected controls were included. Single-cell and peripheral-blood studies were excluded because unavailable for suicide. Curated suicide and BD DEG datasets were integrated with a published bulk transcriptomic dataset of MDD. We identified both shared and disease-specific molecular signatures across suicide, BD, and MDD, and identified DEGs shared across brain regions and those that were region-specific.

A total of 276 genes were shared across all three conditions, implicating cellular stress, hypoxia (*VEGFA, ADM, NDRG1, DDIT4*), immune and inflammatory signaling through TNFα/NF-κB (*IL1B, ICAM1, NFE2L2*) and IL6-JAK-STAT3 pathways (*STAT3, SOCS3, IL1R1m OSMR*), complement activation (*CIR, SERPiNA1, LGALS3*), extracellular matrix remodeling (*COL4A1, SERPINH1, PLOD2*), and unfolded protein response (*BAG3, WFS1, MTHFD2*). Suicide-associated DEGs were prominently downregulated, and involved metal-ion homeostasis, stress responses, synaptic regulation, and circadian pathways, whereas BD showed downregulation in synaptic and vesicle-related processes alongside region-specific metabolic dysregulation. MDD exhibited changes in RNA processing, ribosomal function, and protein synthesis. Genetic integration revealed that 133 genes shared between BD and MDD had gene variants associated with the corresponding condition per GWAS. Transcription factor analysis identified 20 regulatory factors with altered activity across all three conditions despite differences in their predicted downstream targets. Overall, these findings suggest that suicide, MDD, and BD share aspects of their molecular and regulatory pathophysiology while exhibiting distinct patterns, including in a region-specific manner.

## INTRODUCTION

Suicide is a complex and multifactorial phenomenon that remains a major global public health concern. It is the second leading cause of death in the U.S. and the fourth leading cause globally in 15-34-year-olds, with about 800,000 lives lost to suicide annually worldwide^1^. Underlying this mortality is a complex interplay of vulnerabilities, as suicide has genetic^2^ and epigenetic^3^ components and is transdiagnostic, with a psychiatric diagnosis occurring in 70-90% of cases. While major depressive disorder (MDD) is implicated in 60% of suicide cases, other diagnoses include bipolar disorder (BD), alcohol use disorder, substance use disorders, schizophrenia, schizoaffective disorder, and post-traumatic stress disorder, highlighting the complexity of suicide etiology^2^.

MDD is a debilitating psychiatric condition that affects hundreds of millions of people worldwide^3^ imposing substantial burdens on individuals^4^ and society^5^. About 30% of patients are considered treatment-resistant – not responding to at least two adequate treatment regimens^6^ – highlighting the need for new treatments, which can only be achieved with a better understanding of the biological mechanisms underlying the disorder. Family studies estimate the heritability of MDD to be at least 30%^7,8^, including by genome-wide association studies (GWAS)^9^. The identified associated single nucleotide polymorphisms (SNPs)^10–13^ provide important leads for further investigating the genetic and biological basis of MDD.

Though MDD is implicated in most suicide cases, BD carries the highest risk of suicidal behavior among other psychiatric disorders^14–16^, as a significant number of individuals diagnosed with BD may undergo at least one episode of self-harm in their lifetime, while approximately 20% die by suicide^17,18^. BD is a lifelong psychiatric condition characterized by recurrent episodes of mania or hypomania and depression^19^. Though it is estimated that one in about 200 people live with BD globally, many remain misdiagnosed or untreated^20^, as BD can be difficult to distinguish from MDD upon first presentation^21^, and the absence of validated clinical biomarkers leaves clinicians reliant on patient interviews and symptom-based assessment^22^. This subjective diagnostic process contributes to misdiagnosis and delayed treatment, highlighting the necessity of defined molecular signatures of BD to identify reliable biomarkers and novel therapeutic targets.

When investigating the underlying biological mechanisms linking suicide to mood disorder, it is important to focus on human data, as the absence of a direct non-human analogue of suicide presents a limitation to the use of animal models when studying suicide. However, GWAS have shown that gene variants are associated with modest increases in suicide risk^2,23,24^ limiting their predictive value^23^. Higher rates of childhood and recent trauma exposure in suicide decedents – combined with multi-fold higher rates of genome-wide DNA methylation^24–29^ – suggest the need to study the combined effects of genetics and epigenetics for a better understanding of suicide pathogenesis. Postmortem human brain transcriptomics provides an important approach for investigating gene expression profiles associated with mood disorders and suicide at the regional level.

Despite growing evidence of transcriptional dysregulation in MDD, BD, and suicide, the extent to which these conditions share or exhibit distinct molecular signatures remains unclear. Differences in study design, brain regions examined and reported DEGs have limited the identification of reproducible transcriptional changes and shared biological pathways across studies. To address this gap, we systematically integrated bulk-RNA transcriptomics datasets from 37 studies identifying the shared and divergent DEGs and molecular pathways across suicide, MDD, and BD, and analyzing brain-wide and region-specific alterations. This approach can identify dysregulated genes, characterize regional patterns of molecular dysregulation, and determine whether distinct gene-level findings converge on shared biological pathways. Importantly, examining both shared and distinct transcriptomic signatures across conditions may provide a framework for understanding the molecular relationship between mood disorders and suicide, and which are the unique features in people that end up dying by suicide.

## RESULTS

### Retrieved and Included Papers

The literature search using the “RNA sequencing,” “gene expression,” and “brain” identified 37 studies meeting the predefined eligibility criteria Eligible studies examined postmortem human brain tissue, employed untargeted, genome-wide transcriptomic profiling (RNA sequencing or microarray), provided complete lists of differentially expressed genes (DEGs), and directly compared suicide cases with psychiatrically matched non-suicide cases sharing the same underlying diagnosis (e.g., MDD-suicide vs. MDD-non-suicide). This matching strategy was intended to distinguish suicide-associated transcriptional changes from those attributable to the underlying psychiatric disorder. The study selection process is illustrated in the flow diagrams (Supplementary Figure.1 and Supplementary Figure.2). There were four bulk brain transcriptomic studies of suicide^42–45^, 13 of BD^42,46–57^, and 20 of MDD^45,55,58–74^. Collectively, these studies investigated whole-transcriptome gene-expression changes across multiple postmortem brain regions. Detailed characteristics of the included studies are provided in Supplementary eTable 1 (Suicide), eTable 2 (BD), and eTable 3 (MDD).

### Bulk Transcriptomic Signatures in Suicide and Bipolar Disorder

#### Suicide

The four bulk brain transcriptomic studies on suicide revealed a total of 656 suicide-associated DEGs across five brain regions (eTable 4). The number of reported DEGs varied substantially across studies, ranging from six genes in the Pantazatos *et al*. to 330 genes in the Arasappan *et al*. (Fig.1a). Cross-study comparison identified a set of 12 recurrent suicide-associated DEGs across independent studies, with 11 concordantly upregulated and one (*GNA14*) showing discordant direction of regulation in suicide (Fig.1b, Supplementary Figure 3). Sequeira *et al*. and Arasappan *et al*. showed substantial finding overlap, including elevated expression of *ANK2, CD59, GRP51, GNM14, JAK3, LTF, MT1X, MT2A, SEMA4B*, and *SERPINE1* (Fig.1c). In contrast, Mamdani *et al.* shared two upregulated genes, including *CRH and MT2A*. Pantazatos *et al*. showed no gene overlap with other studies. Notably, *MT1X* and *MT2A* were identified across studies, highlighting altered metallothionein-mediated metal homeostasis in suicidal brains.

Across brain regions, the subgenual anterior cingulate cortex (sgACC) had the highest number of DEGs, 328, dominated by 309 downregulated genes and 19 upregulated. The nucleus accumbens (NAc) showed 134 DEGs, including 82 upregulated and 52 downregulated genes, and the anterior cingulate cortex (ACC) had 133 DEGs with 83 upregulated and 50 downregulated. The dorsolateral prefrontal cortex (DLPFC) and the prefrontal cortex (PFC) were reported to have 77 DEGs with 39 downregulated and 17 upregulated and 21 with no reported directional change, while the anterior insula had six DEGs, all of which were downregulated (Fig.1d, eTable 7). The top five up- and downregulated genes for each brain region are shown in Fig.1e. Direction of expression for shared DEGs were mostly consistent across the brain regions, except for *GNA14*, which was upregulated in the DLPFC while downregulated in the sgACC (Fig.1f).

Gene set enrichment analysis of brain-wide suicide-associated DEGs identified significant upregulated pathways related to axon projection and cytoskeletal organization (*SPTBN1, CTNNB1, RAB35, MBP*) (Fig.1g, eTable 9). Conversely, the downregulated pathways were related to metal ion homeostasis and detoxification including metallothionein-mediated metal binding^75^ (*MT1A, MT1E, MT1F, MT1G, MT1H*), wound healing *(ANO6, ANXA2, C1QTNF1, CASP7, CD44)*, and adaptive immune response (*ARID5A, BCL3, C1R, C1RL, C1S, C2*) (Fig.1d).

Region-specific pathway enrichment analysis identified shared enrichment of pathways related to metal-ion homeostasis, inflammatory signaling, vesicular trafficking, and polyamine metabolism. In the sgACC, the most prominent downregulated pathways included wound healing, adaptive immune response, positive regulation of cytokine production, interleukin signaling, and responses to lipopolysaccharide, with *ANXA2*, *BCL3*, *ARID5A*, and *CASP1* among other genes. The most prominent upregulated pathways included response to cocaine, learning and memory, endocrine process, cAMP signaling and Heme biosynthesis, with *ARID5A, BCL3, C1R, C2, AMD,* and *BDNF* among other genes. In ACC, multiple pathways related to metal-ion and copper homeostasis were downregulated, predominantly involving metallothioneins (*MT1G, MT1E, MT1F*) and apoptosis-related genes (*SERPINB13*, *SERPINE1*). In the NAc, downregulated genes were enriched for metallothionein mediated metal binding and responses to metal ions (*MT1H, MT1G, MT1F*), as well as calcium-dependent protein binding involving *ANXA2, S100C,* and *ANXA11*. The anterior insula showed downregulated genes enriched for vesicle/secretory granule pathways and regulation of hydrolase activity, with *SERPINA3* and *CHIL3L1* genes, together with development-related *SLC39A14*. In the DLPFC/PFC, pathways related to circadian cycle, hormone signaling and hydrolase activity were upregulated, including genes such as *CRH, PER3* and *AGMAT*. Downregulated pathways included cellular response to zinc ions, vesicular transport and membrane trafficking, involving *AP1M1, ATP13A2, CD59, MT1A, MT2A, HTR2A*, and GABRA2 (Fig.1h, eTable 10).

#### Bipolar Disorder

A total of 12,970 unique DEGs (eTable 5) associated with BD were identified across six brain regions from 13 independent transcriptomic studies (Fig.1i). Cross-study comparison identified a set of 3,331 recurrent BD-associated DEGs across independent studies, with 1,279 genes consistently downregulated, 1,221 genes consistently upregulated, and 831 discordant genes (Supplementary Figure 4). The top 20 genes shared across the study are shown in Fig.1j. Direction of regulation for shared genes were mostly consistent across the studies, except for genes *ZDHHC11, CDKN1A*, and *AIF1L* (Fig.1k). The number of DEGs varied in between studies and brain regions (Fig.1l), with different studies focusing on different brain regions, including sgACC, ACC, DLPFC/PFC, amygdala, NAc, and hippocampus. The largest dataset was reported by Yan *et al*., which identified 6,865 DEGs in the ACC and 1,351 DEGs in DLPFC/PFC, whereas other studies contributed region-specific gene sets. The proportion of upregulated and downregulated DEGs also differed substantially across the brain regions and the studies.

Region-specific results were heterogeneous (Fig.1l-n). The ACC exhibited the highest number of DEGs with 7,871 genes, followed by 5,772 genes in DLPFC/PFC and 3,859 in amygdala, the most studied brain regions. Comparatively fewer DEGs were detected in sgACC, with 620 genes, and anterior insula – 441 genes, whereas the less studied hippocampus had 25 DEGs and only one was found in NAc. The direction of gene regulation differed across brain regions (Fig. 1l). In the ACC, 4,357 genes were upregulated, while 2,514 genes were downregulated and 22 genes had no established directional change. In the DLPFC/PFC, 2832 genes were upregulated and 2940 were downregulated, with 136 genes reported as both up- and downregulated across studies, and 2,056 upregulated and 1,803 downregulated in the amygdala. Conversely, most genes in the sgACC were downregulated, with 367 downregulated genes versus 253 upregulated genes. The anterior insula contained 294 upregulated genes and 147 downregulated (Fig 1l, eTable 7). Top 10 up- and downregulated genes for each brain regions are shown in Fig.1m. Across the brain regions, direction of regulation for shared DEGs were mostly consistent, except for genes such as *SERPINA3, ABCA6*, *ARID5 and HILPDA* (Fig.1n, eTable 8).

**Figure 1.**
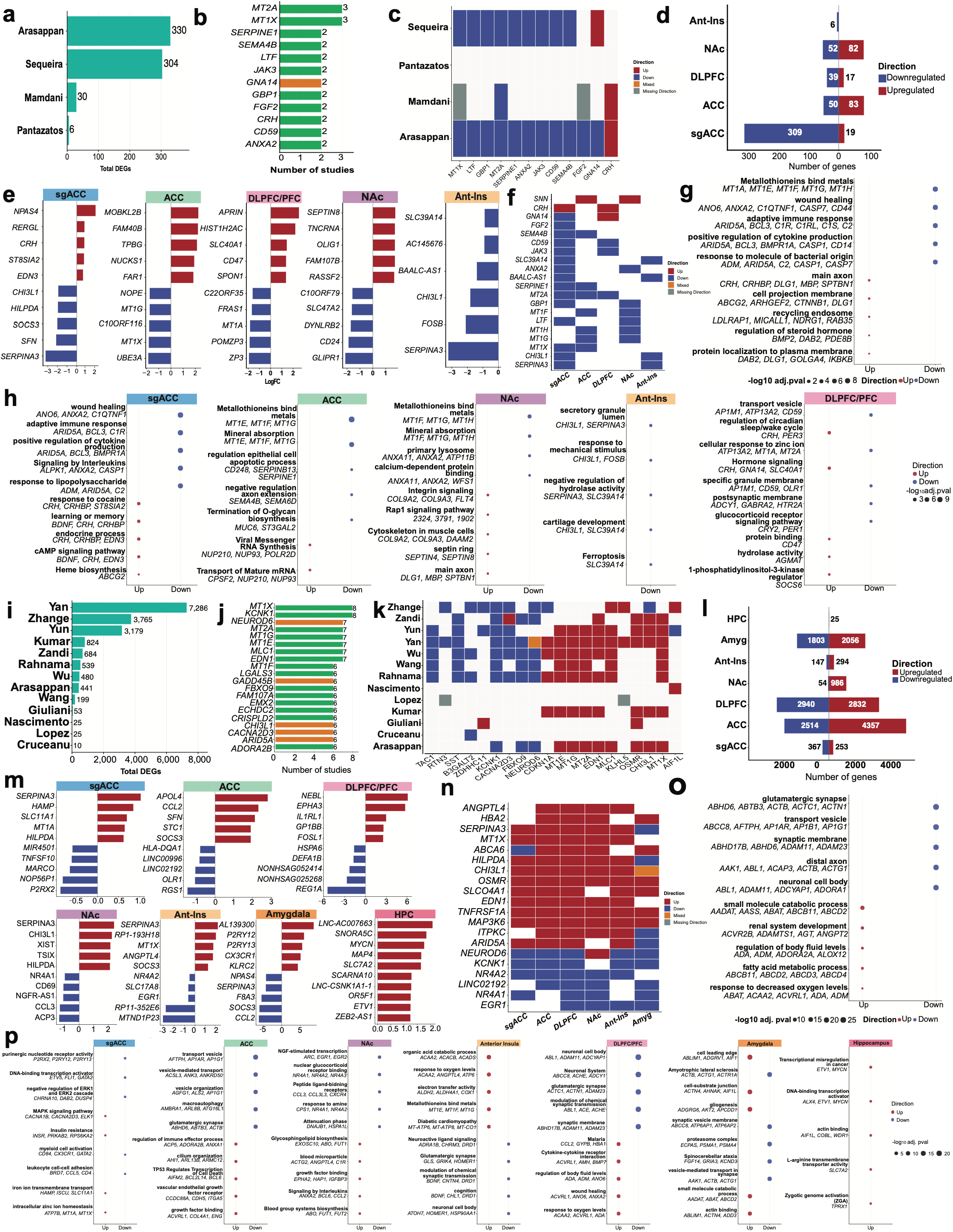
Bulk differential gene expression in suicide and bipolar disorder. **(a)** Total DEGs reported in each suicide study. **(b)** Suicide-associated DEGs shared across ≥ 2 studies, categorized by concordant (green) vs discordant (orange) direction. **(c)** Direction of expression for each shared suicide DEG across individual studies in each study (white = not reported, grey = direction not given/missing, orange = conflicting within a study). **(d)** Number of upregulated (red) and downregulated (blue) suicide DEGs by brain region. **(e)** Top five up- and downregulated suicide DEGs ranked by mean logFC within each brain region. **(f)** Direction of suicide DEGs shared across brain regions. **(g)** Top GO/KEGG/Reactome terms enriched among upregulated vs downregulated suicide DEGs. **(h)** Most significant GO/KEGG/Reactome terms by brain region among suicide DEGs, split by direction. **(i)** Total DEGs reported in each bipolar disorder (BD) study (13 studies). **(j)** BD-associated DEGs shared across ≥ 2 studies, categorized by concordant (green) vs discordant (orange) direction. **(k)** Direction of each shared BD DEG in each study. **(l)** Number of upregulated (red) and downregulated (blue) BD DEGs by brain region. **(m)** Top up- and down-regulated BD DEGs ranked by mean logFC, per region. **(n)** Direction of BD DEGs shared across brain regions. **(o)** Most significant GO/KEGG/Reactome terms enriched among upregulated vs downregulated BD DEGs. **(p)** Top GO/KEGG/Reactome terms by brain region among BD DEGs, shown separately for upregulated and downregulated genes.

Functional enrichment analysis of all BD-associated DEGs revealed pathways involved in synaptic structure and vesicle-mediated transport (Fig.1o, eTable.11). Pathways related to small molecule catabolic process (*AADAT, AASS, ABAT, ABCB11, ABCD2*), regulation of adenosine (*ADA, ADORA2A*), vasculature (*ADM*), inflammation (*ALOX12*), fatty acid metabolic process (*ABCB11, ABCD2, ABCD3, ABCD4*), response to oxygen levels (*ABAT, ACAA2, ACVRL1, ADA, ADM*) were upregulated in BD, whereas pathways related to synaptic membrane (*ABHD17B, ABHD6, ADAM11, ADAM23*) and vesicle transport (*ABCC8, AFTPH, AP1AR, AP1B1, AP1G1*) were downregulated (Fig.1o).

Region-specific pathway enrichment analyses identified distinct pathway profiles across brain regions (Fig.1p, eTable.12). The sgACC had downregulated pathways enriched for purinergic/nucleotide receptor signaling (PRX2, P2RY12, P2RY13), transcriptional regulation (ETV5, DUSP5, DAB2) and immune terms such as myeloid cell activation and leukocyte cell-cell adhesion. Upregulated terms included metal ion homeostasis terms for both iron (*HAMO, ISCU, SLC11A1*) and zinc (*MT1A, MT1X*). The DLPFC/PFC had downregulated genes enriched for neuronal cell body (*ABL1, ADAM11, ADCYAP1*), the neuronal system (*ABCC8, ACHE, ADCY1*), glutamatergic synapse (*ACTC1, ACTN1, ADAM23*), and modulation of chemical synaptic transmission (ABL*1, ACE, ACHE*). Upregulated terms included immune/inflammatory pathways, and wound healing. The hippocampus had upregulated terms including DNA-binding transcription activator activity (*ALX4, ETV1, MYCN*), actin binding (*A1F1L, COBL, WDR1*) and transcriptional dysregulation (*ETV1, MYCN*) (Fig.1p, eTable.12).

### Shared Molecular Features Across Suicide, BD, and MDD

#### Shared Gene Dysregulation Across Conditions and Brain Regions

To identify shared transcriptomic signatures across psychiatric disorders, DEGs found in suicide, BD, and MDD were compared (Fig.2a, eTables.4-6). Comparison of the three datasets identified genes shared across all three conditions as well as genes shared between pairs (eTable 13). The largest overlap was observed between BD and MDD with 6,790 genes, which had the large number of studies, followed by smaller overlaps of 138 genes between suicide and MDD, and 127 genes overlapping between suicide and BD. A total of 276 genes were shared across all three conditions (Fig.2b). The directional expression of these DEGs was compared, and the DEGs were further grouped as i) directionally consistent across all three disorders, ii) consistent between two disorders only, or ii) mixed/other (Fig.2c). Across suicide, BD, and MDD, 12 genes were consistently upregulated, whereas 8 genes were downregulated across all three conditions. Between MDD and BD, there were 62 shared upregulated DEGs and 5 downregulated ones, but they exhibited opposing expression in suicide with 46 genes downregulated in MDD and suicide while upregulated in BD, 13 genes upregulated in suicide and BD but downregulated in MDD and 16 genes downregulated in suicide but upregulated in MDD. The remaining 75 genes did not exhibit a consistent pattern of pairwise concordance (Fig.2c, eTable 14).

**Figure 2.**
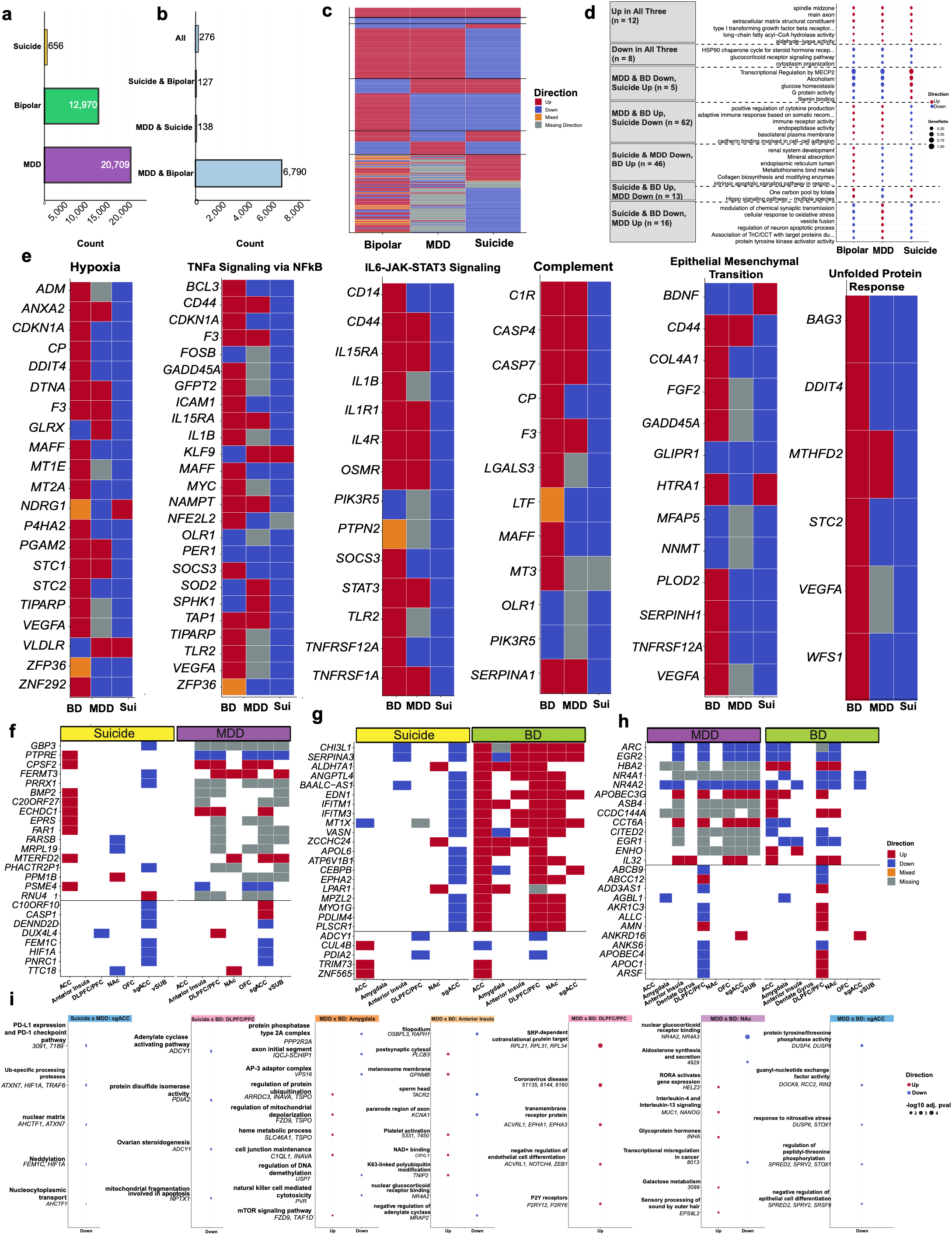
Shared DEGs and pathways across suicide, bipolar disorder (BD), and MDD. **(a)** Total number of bulk DEGs identified for each disease. **(b)** Number of DEGs shared between each pair of diseases and across all three diseases. **(c)** Genes shared across all suicide, BD and MDD, grouped by concordance pattern, showing the direction of each gene in BD, MDD, Suicide. **(d)** Top GO/KEGG/Reactome terms for each concordance pattern in (c), shown separately by direction of expression. **(e)** Direction of genes shared across all three diseases within six Hallmark pathways: Hypoxia, TNFα signaling via NF-κB, IL6-JAK-STAT3 signaling, Complement, Epithelial–mesenchymal transition, Unfolded protein response. **(f)** Suicide and MDD shared genes across regions. **(g)**Suicide and BD shared genes across regions. **(h)** MDD and BD shared genes across regions**. (i)** Significant GO enrichment among pairwise disease overlaps by brain region; only statistically significant enrichments are shown.

#### Shared Altered Pathways Across Conditions and Brain Regions

To characterize shared molecular pathways across conditions, functional enrichment analyses were performed separately for each DEGs with the same expression directionality using Gene Ontology (Biological Process, Molecular Function, and Cellular Component), Reactome, and KEGG, with the top eight enriched terms displayed for each comparison (Fig.2d, eTable.15).

Genes that were concordantly upregulated across all three conditions were enriched for pathways related to spindle and extracellular matrix structural organization, myelin sheath, and aldehyde-lyase activity. Conversely, genes that were concordantly downregulated were enriched for glucocorticoid and corticosteroid receptor signaling, cytoplasm organization, and mesenchymal cell apoptotic process. Genes that were upregulated in suicide and downregulated in both BD and MDD were enriched for pathways related to MECP2-mediated transcriptional regulation, cAMP signaling, alcoholism, transcriptional regulation by NRSF, and NTRK2 signaling. Genes that were downregulated in suicide and MDD but upregulated in BD were enriched for pathways associated with metal ion responses and detoxification, stress responses to metal ions, detoxification of inorganic compounds, and cellular responses to zinc ions. Genes that were downregulated in suicide and BD but upregulated in MDD were enriched for pathways related to synaptic signaling and vesicle trafficking, including regulation of trans-synaptic signaling, modulation of chemical synaptic transmission, regulation of vesicle fusion, and protein tyrosine kinase activator activity (Fig.2d, eTable.15). To further characterize shared molecular signatures, all genes shared across suicide, BD, and MDD were annotated using Hallmark, Reactome, and KEGG pathway databases (eTable.16). Based on their relevance to suicide, BD and MDD pathology, six biological pathways were selected for further study, including hypoxia, TNFα signaling via NF-κB, IL6–JAK– STAT3 signaling, complement, epithelial-mesenchymal transition (EMT), and the unfolded protein response (UPR) (Fig.2e). Across all six pathways, genes were predominantly downregulated in suicide and upregulated in BD, whereas MDD exhibited a more heterogeneous pattern of expression. Hypoxia-associated genes included *HIF3A, VEGFA, DDIT4, P4HA2*, and *CA9*. TNFα signaling via NF-κB included genes such as *TNFAIP3, NFKBIA, PTGS2, ICAM1, CXCL2,* and *IL6ST,* whereas IL6–JAK–STAT3 signaling included *STAT3, IL1R1, IL15RA, TNFRSF1A, and CSF1*. Complement pathway genes included *C1QA, C1QB, C3, CFB*, and *SERPING1*. The epithelial mesenchymal transition pathway comprised of genes associated with extracellular matrix organization and cell adhesion, including *COL4A1, COL4A2, COL5A1, BDNF,* and *GLIPR1*, whereas the unfolded protein response included genes involved in protein folding and endoplasmic reticulum stress, including *CALR, BAG3, HSPA5, HSP90B1, and PDIA4* (Fig.2e).

#### Region-Specific Gene Dysregulation Shared Across Conditions

Region-specific analyses of shared DEGs demonstrated distinct patterns of concordant and discordant regulation across brain regions and disorder pairs. Among concordantly regulated genes, suicide and MDD shared the largest overlap in the sgACC for 13 genes, 8 of which were downregulated in both conditions, including FEM1C, HIF1A, and PNRC1 (Fig.2f, eTable.17). With 10 DEGs in both suicide and BD, the sgACC showed the highest overlap of DEGs in both conditions, though none of them showed same direction of expression in the two diseases (Fig.2g, eTable.18). MDD and BD had the most overlapping DEGs, 938 in the DLPFC/PFC. Of those genes, 120 were upregulated in both diseases, whereas 203 were downregulated, and the directions were not given/missing for remaining (Fig.2h, eTable.19).

#### Region-Specific Altered Pathways Shared Across Conditions

To identify pathways associated with genes altered in the same direction in two conditions and in a specific brain region, GO, KEGG, and Reactome enrichment analysis was performed separately for each region × disease-pair combination with sufficient gene coverage. In the sgACC, downregulated genes in both suicide and MDD were enriched for PDL1 expression, UB-specific processing, nuclear matrix, neddylation and nucleocytoplasmic transport (Fig.2i, eTable.20). In the DLPFC, downregulated genes in suicide and BD were enriched for adenylate cyclase activating pathway, protein disulfide isomerase activity, steroidogenesis and mitochondrial fragmentation (Fig.2i, eTable. 21).

MDD and BD had multi-region significant GO terms. In the amygdala, they had downregulated terms related to protein phosphatase type 2A complex activity (*PPP2R2A*), axon initial segment organization (*IQCJ-SCHIP1*), AP-3 adaptor complex function (*VPS18*), regulation of DNA demethylation (*USP7*), and natural killer cell mediated cytotoxicity (*PVR*), and upregulated terms related to regulation of protein ubiquitination (*ARRDC3, INAVA, TSPO*), regulation of mitochondrial depolarization (*FZD9, TSPO*), heme metabolic process (*SLC46A1, TSPO*), cell junction maintenance (*C1QL1, INAVA*), and the mTOR signaling pathway. In the anterior insula, downregulated pathways included filopodium (*OSBPL3, RAPH1*), sperm head (*TACR2*), paranode region of axon (*KCNA1*), nuclear glucocorticoid receptor binding (*NR4A2*), and negative regulation of adenylate cyclase (*MRAP2*), while upregulated pathways included postsynaptic cytosol (*PLCB3*), melanosome membrane (*GPNMB*), platelet activation, NAD+ binding (*CRYL1*), and K63-linked polyubiquitin modification (*TNIP2*). In the DLPFC, only upregulated terms were observed, including SRP-dependent co-translational protein targeting (*RPL21, RPL31, RPL34*), the coronavirus disease pathway, transmembrane receptor protein (*ACVRL1, EPHA1, EPHA3*), negative regulation of endothelial cell differentiation (*ACVRL1, NOTCH4, ZEB1*), and *P2Y* receptors (*P2RY12, P2RY6*). In the NAc, terms related to interleukin-4/interleukin-13 signaling (*MUC1, NANOG*), glycoprotein hormones (*INHA*), RORA-driven gene expression (*HELZ2*), galactose metabolism, and sensory processing of sound by outer hair cells (*EPS8L2*) were upregulated, while terms involving nuclear glucocorticoid receptor binding (*NR4A2, NR4A3*), aldosterone synthesis and secretion, and transcriptional misregulation in cancer were downregulated. Finally, in the sgACC, only downregulated terms reached significant enrichment, including protein tyrosine/threonine phosphatase activity (*DUSP4, DUSP6*), guanyl-nucleotide exchange factor activity (*DOCK8, RCC2, RIN2*), response to nitrosative stress (*DUSP6, STOX1*), regulation of peptidyl-threonine phosphorylation (*SPRED2, SPRY2, STOX1*), and negative regulation of epithelial cell differentiation (*SPRED2, SPRY2, SRSF6*) (Fig.2i, eTable 22).

### Disease-Specific Dysregulated Genes and Pathways in Suicide, BD, and MDD

#### Disease- and Regional-Specific Gene Dysregulation

To identify molecular alterations unique to each disorder, we studied disease-specific DEGs. MDD showed 13,505 genes (eTable 23), followed by 5,777 genes in BD (eTable 24), and 115 genes were identified as suicide-specific DEGs (eTable 25) (Fig.3a).

Suicide-specific DEGs were prominently identified in the sgACC and ACC, with 39 and 37 respectively. The NAc had 26 DEGs, the DLPFC had 10 and the anterior insula had only one. Interestingly, the DEGs in the sgACC and DLPFC were almost entirely downregulated, with the most downregulated genes including *SMAD1*, RBM27, SERPINE1, and AEBP1 in the sgACC. (Fig.3c).

Among BD-specific DEGs, 2,382 genes were identified in the DLPFC/PFC, 1,550 of which were upregulated – including *FOSL1, LIF,* and *FCGBP* – and 832 were downregulated, including DFA1B, and LCN2 (Fig. 3b,d). In the ACC, 2967 genes were upregulated, and 861 were downregulated; in amygdala, 636 genes were upregulated and 488 were downregulated; and in the sgACC, 98 genes were downregulated and 54 were upregulated. The smallest counts were identified in the in the anterior insula – with 41 downregulated and 104 upregulated genes – and the hippocampus, with only 14 upregulated BD specific DEGs, as there are fewer studies conducted in these regions.

**Figure 3.**
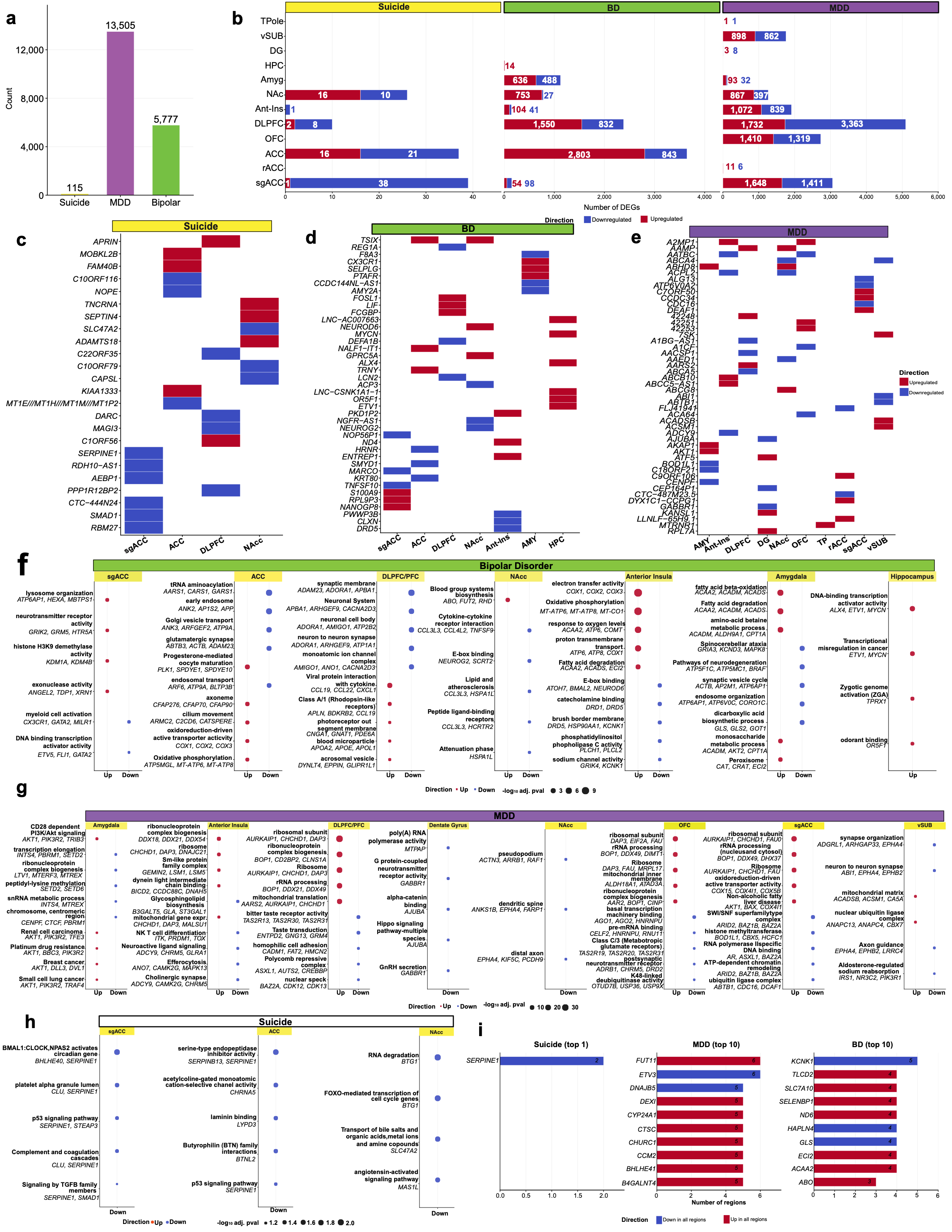
Disease-specific DEG and pathway profiles in suicide, bipolar disorder (BD), and MDD. **(a)** Number of DEGs exclusive to each disease. **(b)** Number of upregulated and downregulated disease-specific DEGs by brain region for each disease. **(c)** Top upregulated and downregulated suicide-specific DEGs by brain region. **(d)** Top upregulated and downregulated BD-specific DEGs by brain region **(e)** Top upregulated and downregulated MDD-specific DEGs by brain region. **(f)** Top BD-specific GO/KEGG/Reactome terms by brain region and direction of expression. **(g)** Top MDD-specific GO/KEGG/Reactome terms by brain region and direction of expression. **(h)** Top Suicide-specific GO/KEGG/Reactome terms by brain region and direction of expression. **(i)** Top 10 disease-specific genes recurring across the most regions with a consistent direction of expression, shown separately for each disease.

MDD-specific DEGs were broadly distributed across brain regions. The DLPFC/PFC contained 5,095 disease-specific DEGs, 3,363 of which were downregulated – including *ABCA5, AACSP1*, and *A1BG-AS1* – and 1,732 were upregulated, including *AARS2*, and *AAMP* (Fig. 3b, e). In the sgACC there were 3,059 DEGs, 2,729 were found in the orbitofrontal cortex (OFC),1,911 in the anterior insula, and 1,264 in the NAc. In contrast, very few disease-specific DEGs were identified in the rostral ACC (rACC), temporal pole, dentate gyrus, and amygdala, much less studied brain regions.

Together, these findings demonstrate distinct regional patterns of transcriptomic dysregulation across disorders, with the DLPFC/PFC appearing as the most studied region in both BD and MDD (Fig.3b).

#### Disease- and Region-Specific Pathway Alterations

Functional enrichment analyses of disease-specific DEGs further highlighted distinct biological processes across disorders.

In BD, the sgACC had upregulated lysosomal, neurotransmitter receptor and chromatin-modifying processes (*ATP6AP1, HEXA, GRIK2, GRM5, KDM1A, KDM4B*) and downregulated immune and transcriptional regulation (*CX3CR1, GATA2, ETV5, FLI1*); the ACC showed downregulated RNA/protein trafficking and synaptic (*AARS1, CARS1, ANK2, APP, ABTB3, ACTB, ADAM23*) and upregulated ciliary and mitochondrial metabolism (*CFAP276, CFAP70, ARMC2, COX1, COX2, COX3*); the DLPFC/PFC had downregulated synaptic/neuronal structure (*ADAM23, ADORA1, APBA1, ARHGEF9, CACNA2D3, ATP1A1*) and upregulated receptor and lipid-transport (*APLN, BDKRB2, CNGA1, GNAT1, APOA2, APOE, APOL1*); the NAc showed downregulated cytokine/receptor signaling and stress-response terms (*NEUROG2, SCRT2, CCL3L3, HCRTR2, HSPA1L*) and upregulated blood group biosynthesis (*ABO, FUT2, RHD*); the anterior insula had upregulated for mitochondria-related oxidative metabolism (*COX1, COX2, COX3, ACAA2, ATP6, ATP8, COMT*) and downregulated neurotransmitter and ion-channel terms (*ATOH7, BMAL2, NEUROD6, DRD1, DRD5, GRIK4, KCNK1*); the amygdala had upregulated fatty-acid/lipid metabolism (*ACAA2, ACADM, ACADS, ALDH9A1, CPT1A*) and downregulated vesicle trafficking and neurodegeneration (*ACTB, AP2M1, ATP6AP1, ATP6V0C, CORO1C*). Notably, the hippocampus showed only upregulated terms that spanned transcriptional and developmental regulation (*ALX4*, *ETV1, MYCN, TPRX1, OR5F1*) (Fig. 3f, eTable26).

In MDD, the amygdala has upregulated PI3K/Akt signaling (*AKT1, PIK3R2, TRIB3*) and downregulated transcriptional/chromatin and RNA-processing (*INTS4, PBRM1, SETD2, SETD6, MTREX, CENPF, CTCF*); the anterior insula showed upregulated ribonucleoprotein/spliceosome assembly and mitochondria gene expression (*DDX18, DDX21, DDX54, CHCHD1, DAP3, GEMIN2, LSM1, BICD2*) and downregulated immune and neuroactive/lipid signaling (*ITK, PRDM1, TOX*); the DLPFC/PFC had upregulated ribosome biogenesis and rRNA processing (*AURKAIP1, CHCHD1, DAP3, BOP1, CD2BP2, DDX21, DDX49, AARS2*) and downregulated cell-adhesion and chromatin remodeling (*CADM1, FAT2, HMCN2, BAZ2A, CDK12, CDK13*). Furthermore, the dentate gyrus had downregulated neurotransmitter-receptor and cell-adhesion signaling (*MTPAP, GABBR1, AJUBA*) and the NAc had downregulated cytoskeletal/dendritic structural (*ACTN3, ARRB1, RAF1, ANKS1B, EPHA4, FARP1, KIF5C, PCDH9*). The OFC had upregulated ribosome biogenesis and mitochondrial terms (*DAP3, EIF2A, FAU, BOP1, DDX49, ALDH18A1, ATAD3A*) and downregulated RNA-binding/transcriptional and neurotransmitter-receptor terms (*AGO1, AGO2, HNRNPU, CELF2, RNU11, ADRB1, CHRM5, DRD2, OTUD7B, USP36, USP9X*); the sgACC showed upregulated ribosome biogenesis and mitochondrial metabolism (*AURKAIP1, CHCHD1, FAU, BOP1, DDX49, DHX37, COX15, COX4I1, COX5B*) and downregulated chromatin-remodeling and ubiquitin-ligase terms (*ARID2, BAZ1B, BAZ2A, BOD1L1, CBX5, HCFC1, AR, ASXL1, ABTB1, CDC16, DCAF1*); and the vSUB had upregulated mitochondrial and ubiquitin-ligase terms (*ACADSB, ACSM1, CA5A, ANAPC13, ANAPC4, CBX7*) and downregulated synaptic function and axon-guidance (*ADGRL1, ARHGAP33, EPHA4, ABI1, EPHB2*) (Fig. 3g, eTable 27).

In suicide, only downregulated terms reached significance across all three regions: the sgACC was enriched for circadian, platelet/coagulation and TGF-β signaling terms (*BHLHE40, SERPINE1, CLU, SMAD1*); protease inhibitor, cholinergic-receptor and cell-adhesion terms (*SERPINB13, SERPINE1, CHRNA5, LYPD3, BTNL2*) were enriched in the ACC; and RNA-degradation, cell-cycle transcription and transport/signaling terms were identified in the NAc (*BTG1, SLC47A2, MAS1L*) (Fig.3h, eTable 28).

### DYSREGULATED GENES WITH RISK VARIANTS PER GWAS STUDIES OF SUICIDE, BD, AND MDD

To determine the extent to which transcriptionally dysregulated genes overlapped with genes that have gene variants found associated with the condition, we next integrated publicly available GWAS data with the DEGs from the 37 studies. This analysis identified 1,223 MDD-, 583 BD-, and 10 suicide-associated DEGs that were also reported as GWAS-associated genes for their respective disorders (Fig. 4a, eTable 29-32).

**Figure 4.**
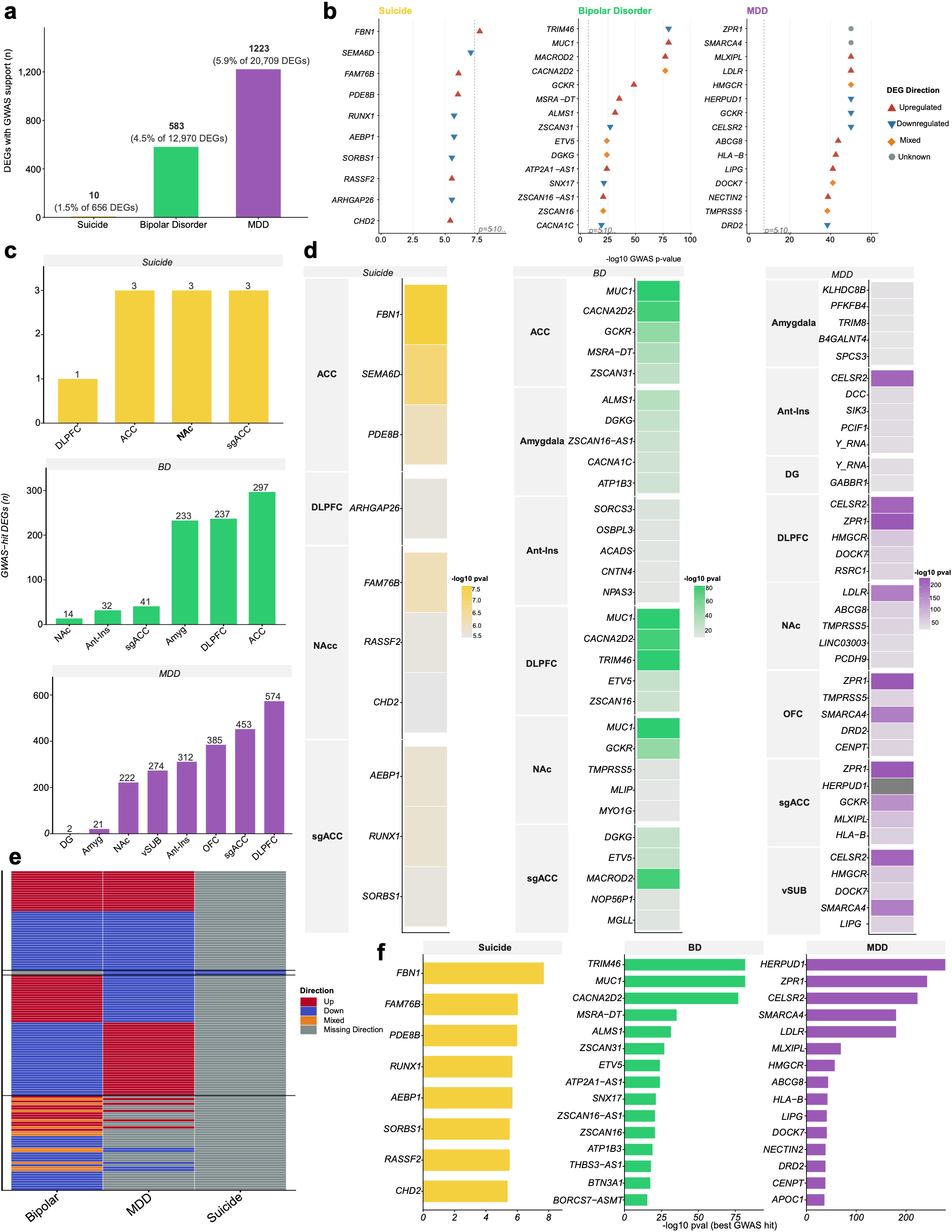
GWAS overlap with total DEGs in suicide, bipolar disorder (BD), and MDD. **(a)** Number and percentage of DEGs overlapped with GWAS hits, for each disease. **(b)** Top 15 GWAS hits ranked by −log10 pvalue) for each disease, with direction of gene expression indicated. **(c)** Number of GWAS-hits DEGs by brain region, for each disease. **(d)** Top 5 GWAS hits by brain region for suicide, bipolar disorder (BD), and MDD. **(e) heatmap for** direction of GWAS-hits DEGs shared across diseases. **(f)** Top 15 GWAS hits for DEGs specific to each disease.

Among the suicide-associated genes, the most significant GWAS hits included *FBN1, SEMA6D, FAM76B,* and *PDE8B*. For BD, the leading GWAS-associated genes included *TRIM46, MUC1, MACROD2,* and *CACNA2D2*. In MDD, the strongest GWAS-associated genes included *HERPUD1, ZPR1, CELSR2,* and *SMARCA4*. Notably, several MDD-associated genes exhibited substantially stronger genetic association signals than the leading genes identified for suicide and BD (Fig.4b). Distribution of GWAS risk genes in each disease and across brain regions are showed in Fig.4c.

GWAS-risk genes showed distinct regional distributions across the three disorders (Fig.4c–d). In suicide, GWAS-risk genes were identified in the ACC, NAc, and sgACC, with three genes identified in each region, while only one gene (*ARHGAP26*) was identified in the DLPFC/PFC. In BD, GWAS risk genes were most abundant in the ACC, with 304 genes, followed by the DLPFC/PFC with 237 genes and amygdala with 233 genes. Comparatively fewer genes were identified in the sgACC, with 41 genes and anterior insula with 32 genes. In MDD, GWAS risk genes were most abundant in the DLPFC/PFC, with 574 genes and sgACC with 453 genes, followed by the OFC with 385 genes and anterior insula with 312 genes. In contrast, only 21 genes were identified in the amygdala and 2 genes in the dentate gyrus. The top five GWAS-risk genes for each brain region across the three disorders are shown in Fig.4d, ranked according to the −log_10_(p-value) of their strongest GWAS association.

Comparison of GWAS-risk genes across disorders revealed substantially greater genetic convergence between BD and MDD than between suicide and either disorder (Fig.4e). MDD and BD shared 133 GWAS hits, including *CACNA1C, ERBB4, NTRK2, TCF4, GCKR,* and *DCC*. In contrast, suicide shared only two GWAS hits with MDD (*ARHGAP26* and *SEMA6D*) and no GWAS hits with BD.

Further study of disease-specific GWAS-risk genes revealed that BD had 450 disease-specific GWAS-risk genes, including *AACS, ABCC1, ACADS, ADAM12,* and *ADCY1*, whereas MDD had 1,088 disease-specific GWAS-risk genes, including *AADAT, ABCA1, ABCG8, ACVR1B,* and *ACO2*. In comparison, suicide had only 8 disease-specific GWAS-risk genes (*AEBP1, CHD2, FAM76B, FBN1, PDE8B, RASSF2, RUNX1,* and *SORBS1*)(Fig.4f, eTable 33).

### Transcription Factor Regulatory Network Analysis Across Suicide, BD, And MDD

To identify putative upstream transcriptional regulators of disorder-specific gene expression signatures, transcription factor (TF) activity was inferred using decoupleR’s implementation of viper against the DoRothEA regulon (confidence levels A–C). Regulon coverage of disease-specific DEGs was higher for BD (677 of 5,777 genes, 11.7%) and suicide (11 of 115 genes, 9.6%) than for MDD (1,007 of 13,505 genes, 7.5%) (Fig.5A, eTable 34). Disorder-specific TF activity analyses highlighted distinct putative regulatory programs. In suicide, *ESR2, ELF5, OTX2*, and *BHLHE22* showed the strongest activation, whereas *EPAS1, NR4A1, ARNTL, FOSL2*, and *FOSL1* showed the strongest repression (Fig.5b). In BD, *STAT2, SP3, HNF4A,* and *ELK1* were strongly activated, while *NCOA1, PAX8, NCOA2*, and *GFI1B* were most repressed (Fig.5c). In MDD, *MYC, POU2F2, MNT*, and *ELK4* showed the greatest activation, whereas *ZNF143, TCF7, E2F5,* and *LEF1* showed the greatest repression (Fig.5d).

**Figure 5.**
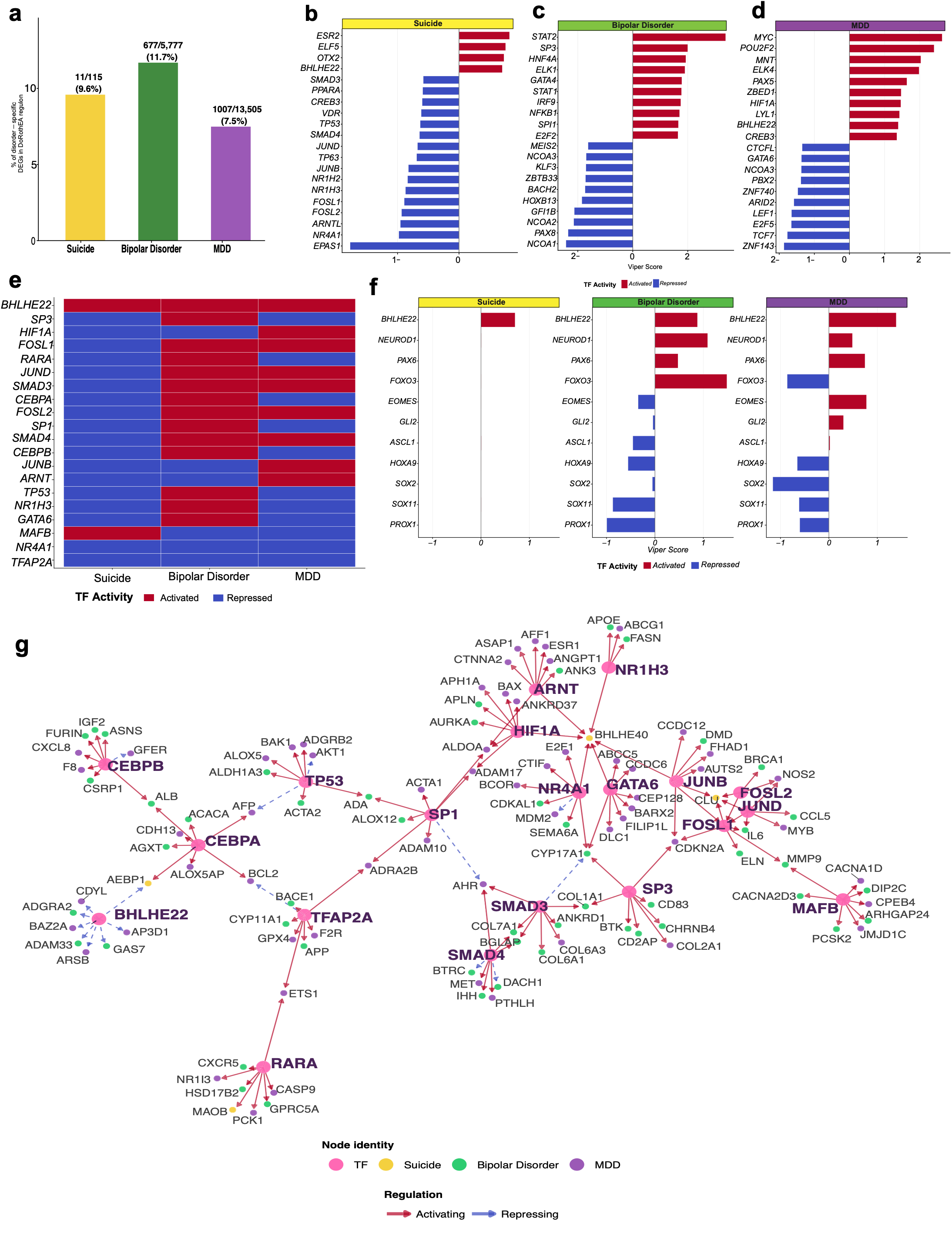
Transcription factor (TF) activity inferred using (VIPER/DoRothEA). **(a)** Percentage of disease-specific DEGs with a high-confidence DoRothEA regulon target annotation. **(b)** Suicide-specific TF activity, top 20 TFs by viper score. **(c)** BD-specific TF activity, top 20 TFs. **(d)** MDD-specific TF activity, top 20 TFs. **(e)** Activity of the 20 TFs with consistent direction across all three diseases. **(f)** TF activity for known immature markers across diseases. **(g)** Regulon network of TFs shared across all three diseases and their target genes, colored by which disease(s) reported each target; arrows show activating vs repressing regulation. * Databases searched: PubMed, Embase, Web of Science. Search terms combined suicide (MeSH term “Suicide” or free text) AND omics/molecular assay terms (RNA sequencing, gene expression, single-nucleus, methylation, proteomics) AND tissue terms (brain or blood) AND a human species filter, joined by Boolean AND across concept blocks. *Source: Page MJ, et al*. *BMJ 2021;372:n71.* doi: 10.1136/bmj.n71. *Licensed under CC BY 4.0*.

Twenty transcription factors were activated/repressed in all three disorders, including *BHLHE22*, which was activated in all three diseases*. FOSL1, RARA, JUND, SMAD3, CEBPA, FOSL2, SP1, SMAD4*, and *CEBPB*, were repressed in suicide but activated in BD while showing a more heterogeneous pattern in MDD. *MAFB* was uniquely activated in suicide while repressed in both BD and MDD (Fig.5e). TFs that are known markers of immature cell types were also explored. Only *BHLHE22* was present across all three disorders, while other markers of immature neuronal cell types were present as TFs in both BD and MDD, including *NEUROD1, PAX6, FOXO3, EOMES, GLI2, ASCL1, HOXA9, SOX2,* and *SOX11* (Fig.5f).

To assess whether the 20 transcription factors with consistent activity across all three conditions converge on a shared regulatory network despite acting on disease-specific genes, we constructed a regulatory network built from these shared transcription factors and their predicted downstream targets, including *CEBPB*, *CEBPA*, *BHLHE22*, *TP53*, *TFAP2A*, *RARA*, *SP1*, *ARNT*, *HIF1A*, *NR1H3*, *GATA6*, *NR4A1*, *SP3*, *SMAD3*, *SMAD4*, *FOSL1*, *FOSL2*, *JUND*, *JUNB*, and *MAFB* (Fig.5e). Because DoRothEA was run separately on each disorder’s DEGs, the predicted targets differed by disorder, though the same 20 transcription factors emerged as regulators across suicide, BD, and MDD.

## DISCUSSION

This study identified both shared and disease-specific molecular signatures across suicide, BD, and MDD. The 276 genes shared across all three conditions point to common dysregulation in cellular stress, hypoxia immune and inflammatory signaling through TNFα/NF-κB and IL6-JAK-STAT3 pathways, complement activation, extracellular matrix remodeling, and the unfolded protein response. Each disease also showed its own profile. Suicide-specific gene signatures were largely downregulated and involved metal-ion homeostasis, stress responses, and synaptic and circadian pathways, whereas BD was downregulated for synaptic and vesicle related processes, alongside region-specific metabolic signatures. MDD exhibited changes in RNA processing, ribosomal function, and protein synthesis. This convergence was also apparent at the level of genetic risk. Genetic integration revealed 133 shared GWAS-associated genes between BD and MDD, further supporting a shared molecular signature between the two disorders. Additionally, transcription factor analysis identified 20 regulators with activity across all three conditions despite differences in their predicted downstream targets. To our knowledge, no previous analysis has comprehensively integrated whole-transcriptome postmortem brain studies to compare shared and disease-specific molecular signatures across suicide, BD, and MDD while accounting for brain-region-specific transcriptional patterns and integrating transcriptomic findings with GWAS risk genes along with transcription factor analysis.

### Suicide: Synaptic Dysfunction, Stress Responses, and Cellular Energy Metabolism

Across four transcriptomic studies, the molecular pathways in suicide related to synaptic dysfunction, altered stress responses, neuroimmune regulation, glial processes, and cellular energy metabolism as recurring features of suicide-associated molecular pathology ^42–45^. This is consistent with previous findings that altered serotonergic and glucocorticoid-related signaling, together with changes in stress-responsive genes, contribute to impaired neuronal adaptation and cellular resilience in MDD-associated suicide^76,77^. Alterations in metallothionein genes further suggest impaired cellular stress responses. In the ACC, six metallothionein genes (*MT1F, MT1G, MT1H,*) were downregulated in suicide, with *MT1F, MT1G,* and *MT1H* also down in the NAc. Because metallothioneins have neuroprotective functions in response to stress and glucocorticoid exposure, their reduced expression may indicate a diminished capacity to protect neurons from chronic stress-related cellular damage^44^. Upregulation of *NPAS4* suggests altered activity-dependent synaptic plasticity and excitatory–inhibitory balance^78^, while *SEPTIN8* reflect changes in synaptic vesicle trafficking and neurotransmitter release. *SEPTIN4*, a presynaptic scaffold in dopaminergic terminals, further implicates altered dopaminergic synaptic function and α-synuclein related neuronal stress. These findings suggest disrupted synaptic plasticity, presynaptic function, and neuronal circuit homeostasis in suicide. *SERPINA3* encodes α1-antichymotrypsin, an acute-phase protein involved in serine protease inhibition and inflammatory regulation. Elevated expression is found in brain tissue and amyloid plaques linked to Alzheimer’s disease^79^

The observed molecular heterogeneity observed in the suicide data is further supported by peripheral and central transcriptomic findings. Four genes showed significant changes in both blood and brain samples between MDD subjects who died by suicide (MDD-S) and those who did not (MDD-NS). *SOX9* was decreased and *PER3* increased in MDD-S in both tissues, whereas *CD19* and *TERF1* showed opposite changes between blood and DLPFC, highlighting both shared and tissue-specific molecular signatures^43^.

Importantly, the reviewed studies also provide evidence for altered cellular energetics in suicide. Pantazatos *et al*. reported alterations in ATPase-related activity^45^, while Mamdani *et al*. identified the mitochondrial carrier gene *SLC25A26* among potential blood biomarkers of suicide in MDD^43^. Considering the enrichment of synaptic and metabolic pathways observed in our analysis, these findings suggest that disrupted cellular energy regulation may contribute to neuronal and synaptic dysfunction. Given the high energetic demands of synaptic transmission, impaired mitochondrial function could compromise ATP-dependent processes required for neurotransmission, cellular homeostasis, and adaptation to chronic stress.

### Bipolar Disorder: Synaptic and Vesicular Dysfunction Alongside Vascular, Immune, and Extracellular Alterations

The strongest functional enrichment in BD involved synaptic and vesicle-related pathways, with downregulation of genes including *RPH3A, PENK, BDNF, and PRKCB,* which are involved in synaptic transmission and neuronal signaling. Similarly, reduced expression of *SYT11, SYT12, MCTP2, UNC13A, MAL2,* and *RAB27B* highlights altered synaptic vesicle trafficking and neurotransmitter release, suggesting impaired neuronal communication as an important feature of BD. Several genes associated with vascular, immune, and extracellular processes were upregulated: *CX3CR1, LRP5, PDGFRA,* and *CYBB* implicate vascular morphogenesis and immune/glial signaling, while *VMA2, OMD, ADAMTS9, TLR3,* and *PODN* implicate extracellular matrix remodeling and cellular interactions. These findings suggest that BD-associated transcriptional changes extend beyond neuronal signaling and may involve coordinated alterations in neuronal, glial, vascular, and extracellular compartments. In addition, BD is characterized by a core signature of altered synaptic and vesicle function, accompanied by region-specific metabolic, vascular, immune, and structural abnormalities. Notably, the involvement of *ACAA2, ACACB, ACADS, ALDH1L1*, and *ATP6* in the anterior insula provides response to oxygen levels and show a potential connection between BD-associated altered cellular energy metabolism, consistent with mitochondrial dysfunction hypotheses of BD^80^

### Convergent and Divergent Transcriptomic Pathways Across Suicide, BD, and MDD

The identification of 276 genes shared across suicide, BD, and MDD indicates substantial molecular convergence across psychiatric disorders. Shared genes implicated hypoxia (*HIF3A, VEGFA, DDIT4, CA9*), inflammatory signaling (*TNFAIP3, NFKBIA, IL6ST*), complement (*C1QA, C1QB, C3, CFB*), extracellular matrix remodeling (*COL4A1, COL4A2, COL5A1*), and unfolded protein response (*CALR, HSPA5, HSP90B1*), suggesting common alterations in cellular stress, immune, and tissue-remodeling processes.

The 62 genes upregulated in BD and MDD but downregulated in suicide were enriched for immune-and membrane-related pathways, such as immune receptor activity, cytokine receptor activity, endopeptidase activity, basolateral and basal plasma membrane components, and cadherin binding involved in cell-cell adhesion. This opposite direction of regulation suggests that suicide diverges from BD and MDD in how these shared immune and membrane-related pathways are controlled, rather than following the same regulatory pattern.

Suicide-specific DEGs marked with downregulation of circadian regulation, dopaminergic signaling, and immune/platelet pathways, including *BHLHE40, SERPINE1, CHRNA5,* and *CLU.* This is consistent with recent genetic evidence linking circadian and sleep-related traits to suicide risk^81^. BD-specific DEGs were more broadly distributed and showed prominent upregulation of metabolic and energy-related pathways. Mitochondrial-encoded genes *COX1–3, CYTB,* and *ND1–4* were upregulated in the anterior insula, while *ACAA2, ACADM, ACADS,* and *AKT2* were upregulated in the amygdala, implicating oxidative metabolism and fatty-acid β-oxidation in BD-specific molecular alterations. This aligns with recent multi-omics evidence showing that energy-metabolism disturbance is more pronounced in BD than in MDD^82^, further supporting disrupted energy metabolism as a core feature of BD-associated molecular pathology.

MDD showed a distinct and more consistent signature of increased protein synthesis and RNA-processing machinery, with upregulation of ribosomal genes such as *RPL10A, RPL13A, RPL14, RPL32,* and *RPL38,* as well as splicing-related genes *CWF19L2, DDX41,* and *LSM5* across the anterior insula and sgACC. This upregulation contrasts with prior postmortem work reporting downregulated ribosomal and protein synthesis genes in MDD in overlapping regions, including the anterior insula and sgACC^42^ and the DLPFC/ACC^83^ suggesting that the direction of ribosomal gene dysregulation in MDD may vary across cohorts, illness stage, or other unmeasured clinical factors rather than reflecting a single consistent pattern. Notably, the DLPFC/PFC showed the largest number of disease-specific DEGs in both BD and MDD, although the direction of regulation differed between disorders. This likely reflects, at least in part, the extensive transcriptomic coverage of this highly studied region and should therefore be interpreted cautiously.

### GWAS-Transcriptome Integration: Shared and Disease-Specific Genetic Risk Across BD, MDD, and Suicide

Integration of GWAS and transcriptomic data revealed both shared and disorder-specific genetic-transcriptional signatures. The identification of 133 GWAS-risk genes shared between BD and MDD, including *CACNA1C*, *ERBB4*, *NTRK2*, *TCF4*, *GCKR*, and *DCC*, suggests shared genetic vulnerability underlying transcriptional dysregulation. In contrast, suicide showed limited genetic overlap, sharing only *ARHGAP26* and *SEMA6D* with MDD and no GWAS-risk genes with BD, potentially reflecting the more limited GWAS data currently available for suicide. Among disease-specific GWAS-risk genes, *ACADS* and *ADCY1* in BD and *ACO2* and *ACVR1B* in MDD highlight potential metabolic and cellular signaling mechanisms specific to each disorder.

### Transcription Factor Convergence with Condition-Specific Downstream Targets

Twenty TFs were shared across all three conditions, suggesting convergence at the level of upstream regulatory mechanisms despite differences in downstream gene expression. *BHLHE22* showed consistent activation across disorders, while specific factors (*FOSL1, FOSL2, JUND, SMAD3, SMAD4, CEBPA, CEBPB,* and *RARA*) showed similar activity patterns in BD and suicide but more heterogeneous activity in MDD. *BHLHE22* repressed all its downstream targets, but the targets themselves differed by disorder: in BD, *ADAM33, ADGRA2, GAS7, PLCL2, and PTPRJ; in MDD, AP3D1, ARSB, BAZ2A, CDYL, IGF1R, SH3KBP1,* and SPON2; and in suicide, only *AEBP1*.

These disease-specific targets had distinct functions per disorder: BD targets clustered around vascular/angiogenic signaling (*ADGRA2, PTPRJ*) and neuronal migration and morphogenesis (*GAS7*); MDD targets clustered around chromatin regulation (*CDYL, BAZ2A*) and lysosomal/vesicular trafficking (*AP3D1, ARSB, SH3KBP1*), whereas suicide’s single target is a transcriptional regulator (*AEBP1*) (Gene Cards). These results illustrate how a single shared upstream regulator can act on functionally distinct downstream programs across disorders (Fig. 5g, eTable 34). These findings support a model in which psychiatric disorders share upstream regulatory and genetic vulnerability but exhibit disorder-specific transcriptional manifestations.

### Limitations

A key limitation of this study is that postmortem tissue does not reveal the mood state of BD patients at the time of death, whether manic, depressed, or euthymic. Future studies incorporating mood state at death would allow for direct comparison of energy metabolism shifts across affective states. This analysis was also restricted to RNA expression due to limited proteomic studies in suicide; further proteomic profiling would add complementary insight since RNA levels do not also reflect protein abundance. Finally, because bulk RNA sequencing averages expression across all cell types in a sample, it cannot resolve molecular changes at the cellular or spatial level.

### Conclusion

Overall, this analysis identifies both shared and distinct transcriptomic signatures across suicide, BD, and MDD, highlighting alterations in synaptic function, cellular stress responses, immune regulation, and energy metabolism. In suicide-associated studies, changes in *MT1F*, *MT1G*, *MT1H*, *NPAS4*, *SEPTIN8*, and *SEPTIN4* suggest potential disruptions in cellular stress protection, synaptic plasticity, and neurotransmission. In BD, alterations in synaptic vesicle-related genes, including *RPH3A*, *SYT11*, *SYT12*, and *UNC13A*, alongside mitochondrial genes such as *COX1–COX3* and *ND1–ND4*, implicate neuronal communication and cellular energy metabolism. In MDD, region-dependent changes in ribosomal function, RNA processing, and protein synthesis suggest alterations in the molecular machinery regulating gene expression. Across all three conditions, shared signatures involving *HIF3A*, *VEGFA*, *TNFAIP3*, *C3*, and *HSPA5* highlight potential convergence in hypoxia responses, immune signaling, and cellular stress pathways. However, differences in expression direction and regional distribution indicate that these processes may be regulated differently across disorders. Overall, these findings provide a framework for understanding convergent and disorder-specific molecular alterations, while further cell-type-resolved and functional studies are needed to establish their biological significance and clarify their relevance to suicide risk.

## METHODS

A systematic literature search was conducted across three databases, PubMed, Embase, and Web of Science, using search terms combining suicide, gene expression, RNA sequencing, transcriptomics, and brain or blood tissue, restricted to human studies. PubMed searches were performed programmatically in R using the easyPubMed package, whereas equivalent search strategies were conducted manually in Embase and Web of Science before export into R for downstream processing. In suicide, four studies were retained for inclusion following screening criteria requiring that studies: (i) examined postmortem brain from human subjects, (ii) employed untargeted, genome-wide transcriptomic profiling (RNA sequencing or microarray), (iii) provided whole DEG lists, and (iv) directly compared suicide cases against psychiatrically matched non-suicide cases sharing the same underlying diagnosis (e.g. MDD-suicide vs. MDD-non-suicide), ensuring that differential gene expression could be attributed specifically to suicide rather than to psychiatric illness. A parallel search using the same pipeline was conducted for bipolar disorder (BD), replacing the suicide MeSH term with “bipolar disorder”[Mesh], and applying equivalent screening criteria to identify BD transcriptomic DEG lists from postmortem human brain. MDD transcriptomic DEG lists were sourced from a prior systematic literature review^84^ and supplemented with RNA sequencing from MDD brain hippocampus^85^.

Disease-specific DEG signatures were then identified by comparing the three DEG lists, suicide, BD, and MDD, and extracting genes unique to each condition using gene-level overlap analysis.

Over-representation analysis was run separately on up- and down-regulated shared and disease-specific gene sets against Gene Ontology terms using clusterProfiler (Benjamini–Hochberg adjusted p < 0.05), with redundant terms collapsed by semantic similarity. ORA was used instead of GSEA because logFC was not comparable across studies and platforms. To evaluate genetic support for transcriptomic findings, DEGs for each disease were integrated with publicly available genome-wide association study (GWAS) summary statistics for suicide, MDD, and BD; genes supported by both differential expression and GWAS evidence within the corresponding disorder were prioritized for downstream interpretation. GWAS-risk genes were also compared directly across disorders to identify genes shared between disorder pairs independent of transcriptomic overlap. In addition disease-specific DEGs were intersected with each disorder GWAS-risk genes to identify those that were unique to a single disorder. Transcription factor regulatory network analyses were performed using decoupleR with DoRothEA regulons to identify putative upstream regulators associated with shared and disorder-specific molecular signatures; downstream target genes in these networks were restricted to disease-specific DEGs, so that shared regulators were evaluated against non-overlapping, disorder-specific transcriptional programs. All statistical analyses, data processing, and figure generation were performed in R using packages from the tidyverse and Bioconductor ecosystems.

## DATA AVAILABILITY

The data associated with the results of this paper are in Supplementary Tables

---

**Supplementary eFigure 1:**
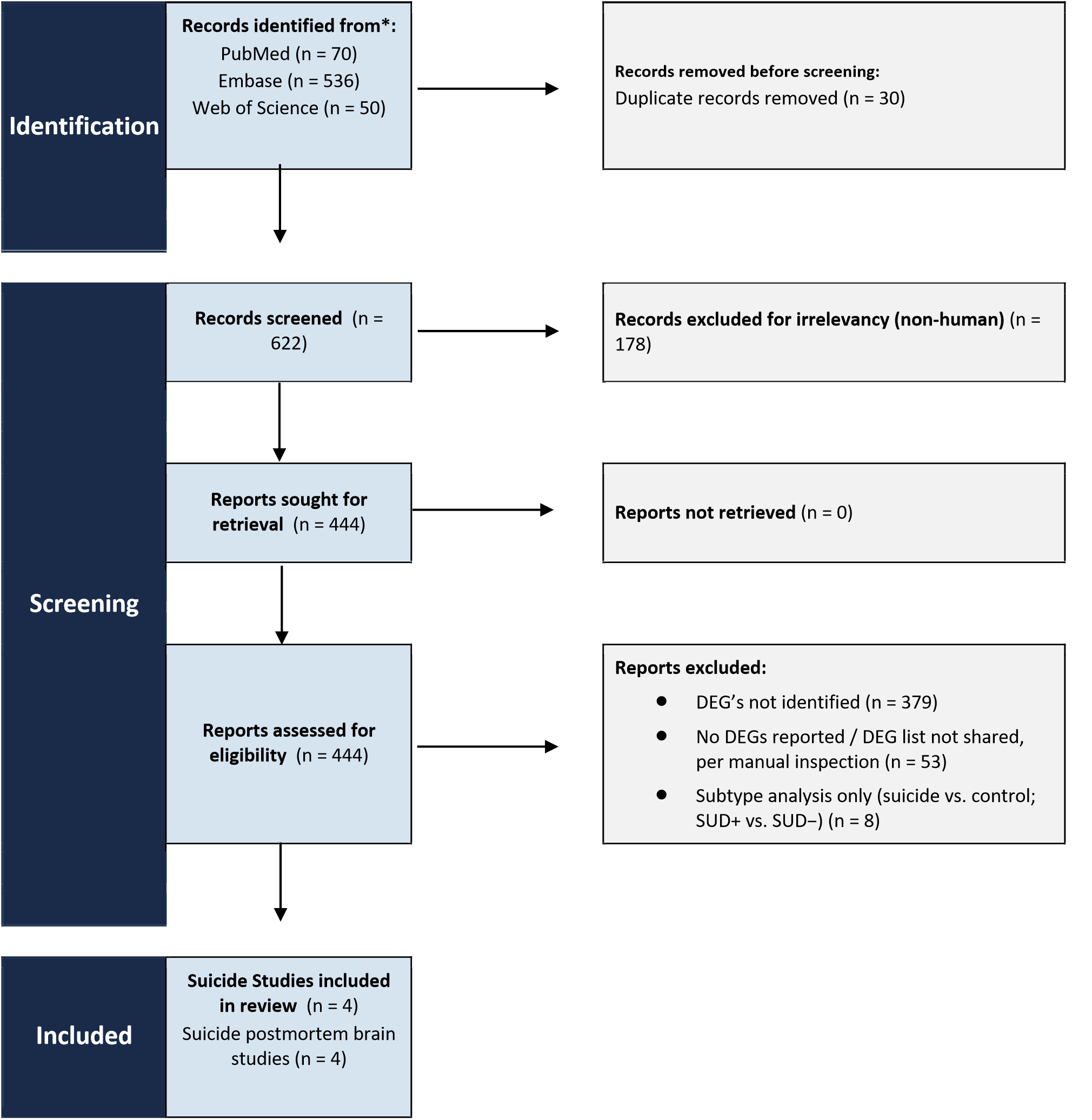
PRISMA Flow Diagram for Suicide. This diagram shows the systematic process we followed to include papers captured by our search. * Databases searched: PubMed, Embase, Web of Science. Search terms combined bipolar disorder (MeSH term “Bipolar Disorder” or free text) AND omics/molecular assay terms (RNA sequencing, gene expression, single-nucleus, methylation, proteomics) AND tissue terms (brain or blood) AND a human species filter, joined by Boolean AND across concept blocks. *Source: Page MJ, et al. BMJ 2021;372:n71.* doi: 10.1136/bmj.n71. *Licensed under CC BY 4.0*.

**Supplementary eFigure 2):**
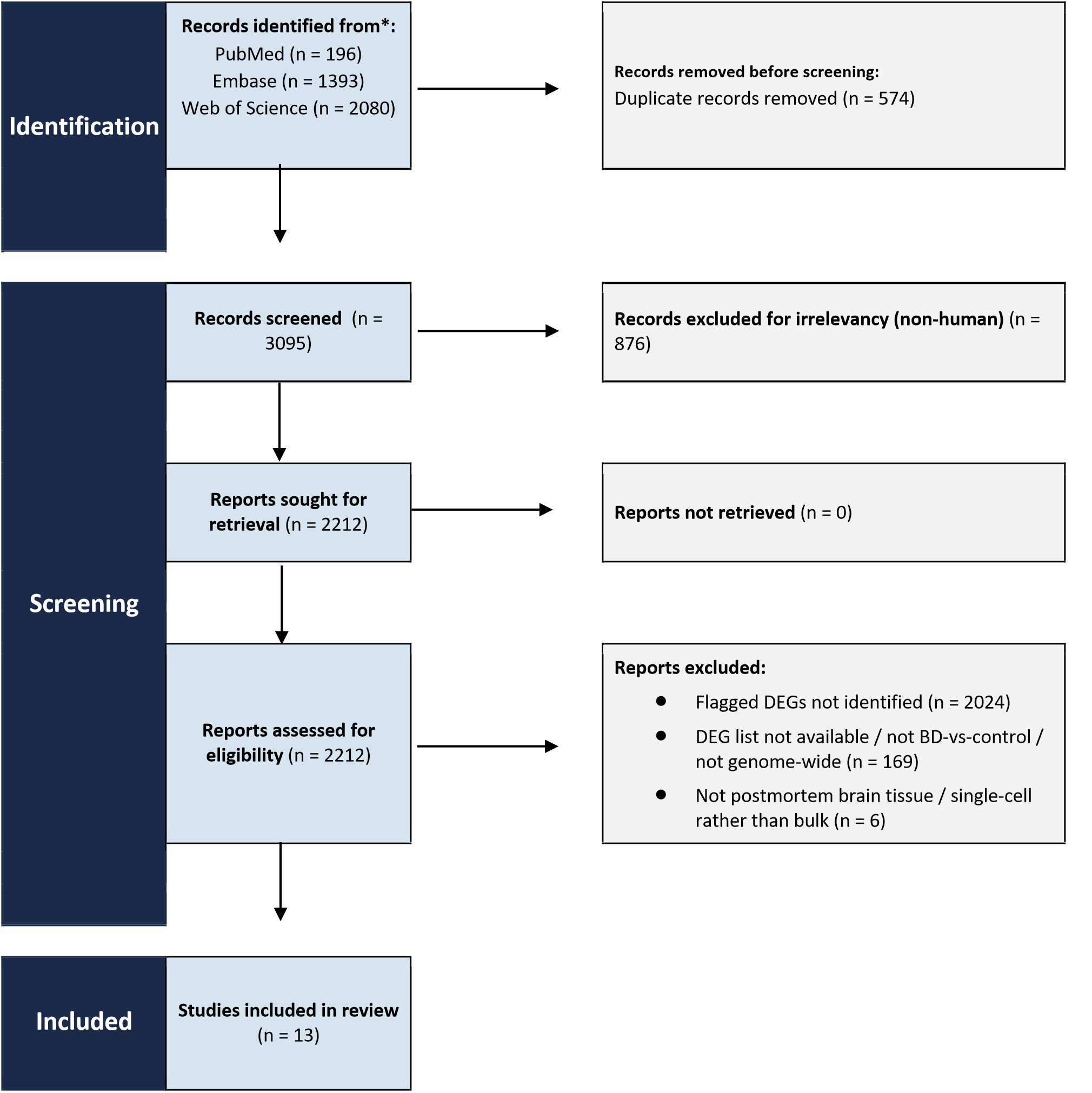
PRISMA Flow Diagram for Bipolar. This diagram shows the systematic process we followed to include papers captured by our search.

**Supplementary eFigure 3:**
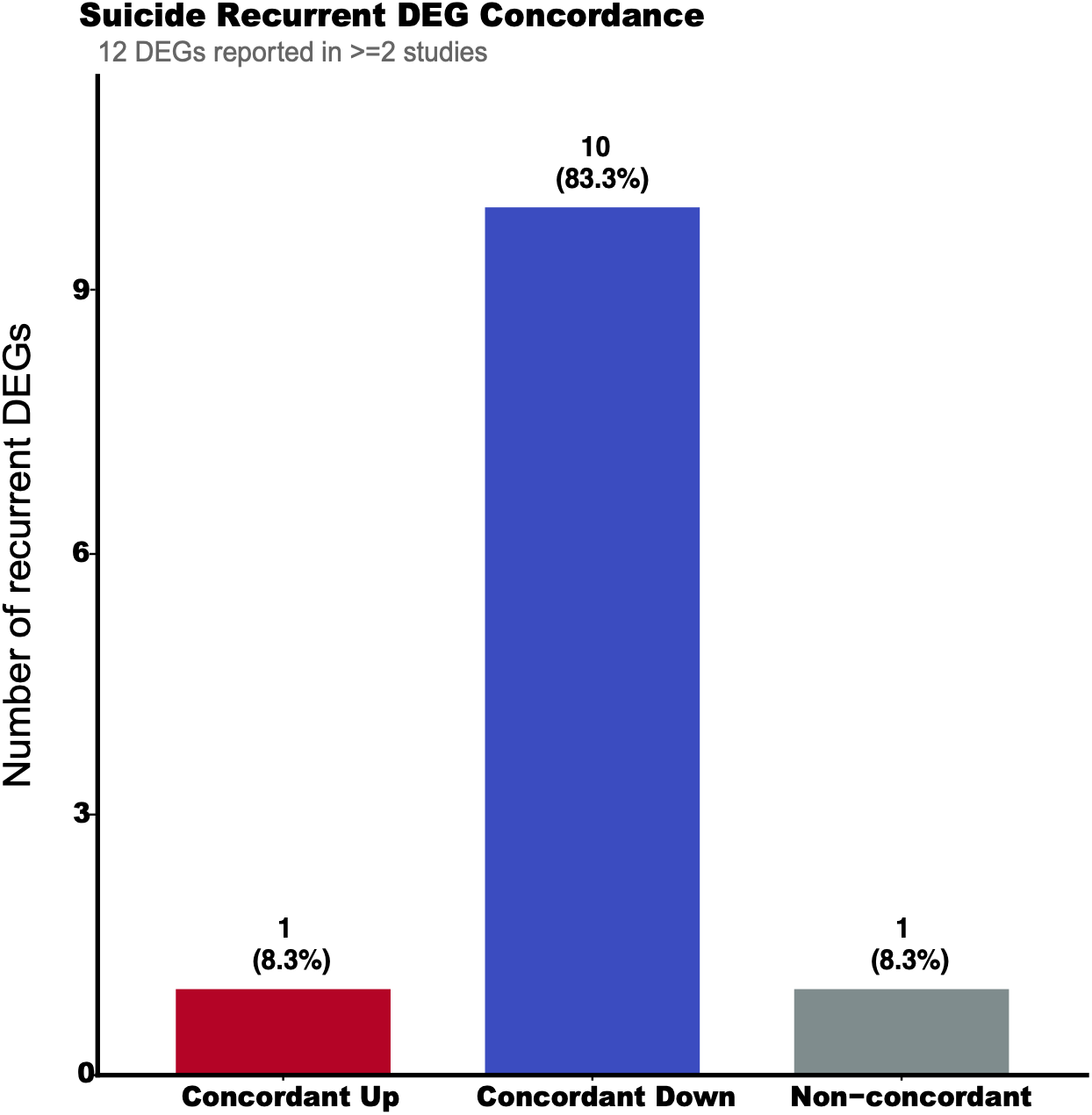
Concordance of recurrent suicide DEGs. The 12 DEGs reported in at least two studies, grouped by whether studies agreed on direction: concordant up, concordant down, or non-concordant (reported as both up and down).

**Supplementary eFigure 4:**
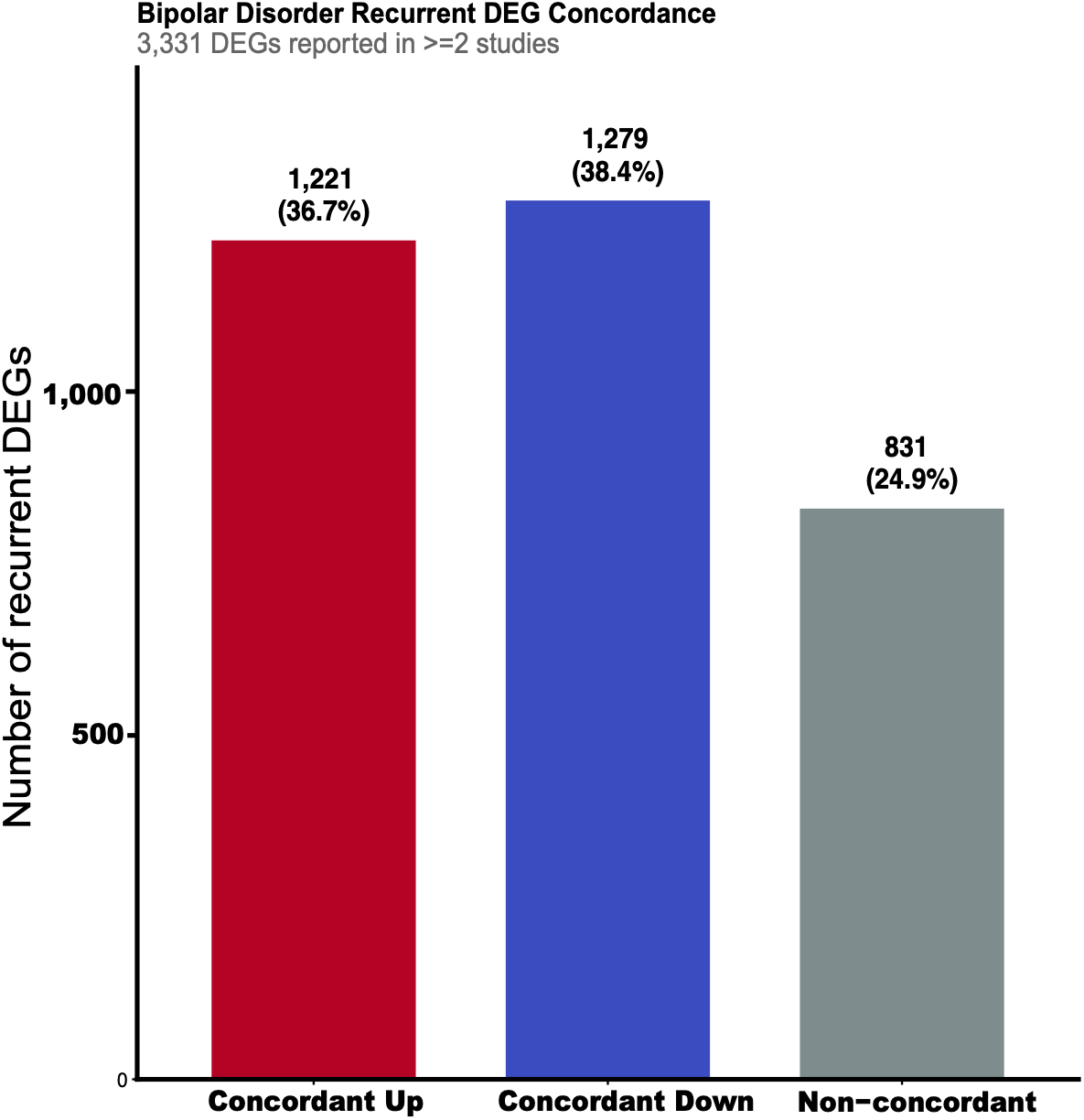
Concordance of recurrent BD DEGs. The 3,331 DEGs reported in at least two studies, grouped by whether studies agreed on direction: concordant up, concordant down, or non-concordant (reported as both up and down).

**eTable 1**: Transcriptomic studies of suicide included in the systematic review (author, method, sample, DEG counts, brain regions)

**eTable 2:** Transcriptomic studies of bipolar disorder (BD) included in the systematic review.

**eTable 3:** Transcriptomic studies of major depressive disorder (MDD) included in the systematic review.

**eTable 4**: Full list of suicide differentially expressed genes (DEGs), with direction, brain region, and source study.

**eTable 5**: Full list of BD DEGs, with direction, brain region, logFC, and source study.

**eTable 6**: Full list of MDD DEGs, with direction, brain region, and source study.

**eTable 7**: Counts of upregulated, downregulated, mixed, and unknown-direction DEGs per disease, by brain region.

**eTable 8**: Per-gene consistency of regulation direction across brain regions, by disease.

**eTable 9**: GO/ORA results for the full suicide bulk DEG list.

**eTable 10**: GO/pathway enrichment for suicide DEGs, stratified by brain region.

**eTable 11**: GO/ORA results for the full BD bulk DEG list.

**eTable 12**: GO/pathway enrichment for BD DEGs, stratified by brain region.

**eTable 13**: Gene counts for each disease-overlap category (disease-specific, pairwise, and shared across all three).

**eTable 14**: Per-gene regulation direction across suicide, BD, and MDD, with concordance classification.

**eTable 15**: GO/pathway enrichment for gene sets showing concordant (same-direction) regulation across disorders.

**eTable 16**: Master gene-level table combining direction, overlap category, gene function/pathways, GWAS support, and TF regulation across all three disorders.

**eTable 17**: Region-level counts of genes shared between suicide and MDD, broken down by direction-of-change combination.

**eTable 18**: Region-level counts of genes shared between suicide and BD, broken down by direction-of-change combination.

**eTable 19**: Region-level counts of genes shared between MDD and BD, broken down by direction-of-change combination.

**eTable 20**: GO/KEGG/Reactome enrichment for genes shared between suicide and MDD, stratified by region and direction.

**eTable 21**: GO/KEGG/Reactome enrichment for genes shared between suicide and BD, stratified by region and direction.

**eTable 22**: GO/KEGG/Reactome enrichment for genes shared between MDD and BD, stratified by region and direction.

**eTable 23**: MDD-exclusive DEGs, with recurrence across studies and brain regions.

**eTable 24**: BD-exclusive DEGs, with recurrence across studies and brain regions.

**eTable 25**: Suicide-exclusive DEGs, with recurrence across studies and brain regions.

**eTable 26**: GO/pathway enrichment for BD-specific DEGs, stratified by brain region.

**eTable 27**: GO/pathway enrichment for MDD-specific DEGs, stratified by brain region.

**eTable 28**: GO/pathway enrichment for suicide-specific DEGs, stratified by brain region.

**eTable 29**: Summary counts of DEGs with GWAS support, per disorder.

**eTable 30**: Suicide DEGs with GWAS support, including trait, p-value, and direction.

**eTable 31**: BD DEGs with GWAS support, including trait, p-value, and direction.

**eTable 32**: MDD DEGs with GWAS support, including trait, p-value, and direction.

**eTable 33**: Disease-exclusive DEGs with GWAS support, by disease and brain region.

**eTable 34**: Full transcription factor-to-target regulatory network (shared TFs), with target gene, regulation direction, confidence, and disease.

